# *De novo* genome assembly of the Tobacco Hornworm moth (*Manduca sexta*)

**DOI:** 10.1101/2020.08.29.270983

**Authors:** Ariel Gershman, Tatiana Gelaf Romer, Yunfan Fan, Roham Razaghi, Wendy A. Smith, Winston Timp

## Abstract

The Tobacco hornworm, *Manduca sexta,* is a lepidopteran insect that is used extensively as a model system for studying insect biology, development, neuroscience and immunity. However, current studies rely on the highly fragmented reference genome Msex_1.0, which was created using now-outdated technologies and is hindered by a variety of deficiencies and inaccuracies. We present the new reference genome for *M. sexta*, JHU_Msex_v1.0, applying a combination of modern technologies in a *de novo* assembly to increase continuity, accuracy, and completeness. The assembly is 470 Mb and is ~20x more continuous than the original assembly, with scaffold N50 >14 Mb. We annotated the assembly by lifting over existing annotations and supplementing with additional supporting RNA-based data for a total of 25,256 genes. The new reference assembly is accessible in annotated form for public use. We demonstrate that improved continuity of the *M. sexta* genome improves resequencing studies and benefits future research on *M. sexta* as a model organism.

## Introduction

As a large, easily grown insect, the tobacco hornworm (*M. sexta, NCBI:txid7130)* is a key model for studying biochemical mechanisms in insect biochemistry, physiology, neurobiology, development, and immunity [1,2,3,4]. In particular, its innate immune pathways, many of which share mammalian homologs, have been characterized in detail and used as a model system for studying fungal virulence and drug efficacy [5,6]. *M. sexta* metamorphosis has also proved an excellent system for studying development, particularly the complex interactions between hormones and environmental regulators, e.g. nutrition, that regulate major developmental changes. More study of this insect has practical applications; *M. sexta* is a prevalent agricultural pest that feeds on solanaceous plants, e.g. tobacco, and has the unique ability to tolerate large amounts of solanaceous alkaloids, such as nicotine [7]. Recent advancements in genomic sequencing have led to chromosome level reference genomes of other lepidopterans such as the domestic silkworm *(Bombyx mori)*, another member of the Bombycoidea superfamily, and the monarch butterfly (*Danaus plexippus*) *[8,9].*

The first effort to sequence and assemble the *M. sexta* genome was an incredible collaboration by experts at 57 different institutions resulting in the Msex_1.0 genome assembly, a 419.4 Mb genome made up of 25.8% repetitive sequence. The accompanying gene set comprised 15,451 protein-coding genes, of which 2,498 were manually curated [10]. However, sequencing technology has advanced significantly since the initial genome release in 2016. The original Msex_1.0 genome comprises 20,869 scaffolds with the largest scaffold being just 3.2 Mb (N50 = 664 Kb) which lags behind the quality of more recently assembled lepidopteran reference genomes such as the Domestic silkworm (*B. Mori),* a 460 Mb genome with a scaffold N50 of 16.8 Mb [8]. Because *M. sexta* is a model species in lepidoptera, its genome assembly needs to be highly accurate and contiguous to allow for comparative genomics studies of insect diversity, biochemical studies on homologous mammalian pathways and agricultural studies regarding the tomato and tobacco crops, a food source for *M. sexta.*

Here, we present the results of a new genome analysis of *M. sexta* based on deep sequencing of short and long reads. We generated an assembly that consists of 470 Mb and a scaffold N50 of 14.2 Mb. We also lifted over the original genome annotation to our new assembly as well as provide high quality novel gene models constructed by combining the new genome assembly with available RNA sequence data. We demonstrate the utility of our new genome assembly through comparative genomics and gene expression analysis throughout *M. sexta* development cementing our assembly as a core resource for the lepidopteran community.

## Materials and Methods

### Sample collection and sequencing

Six moths were purchased as pupae from Carolina Biological (<u>Moths</u>); two of which survived to adulthood. On the day of emergence, moths were euthanized by placing them at 4°C for 10 minutes. The moth’s body, legs, head and wings were snap-frozen in liquid nitrogen, then stored at −80°C prior to DNA extraction. High-quality genomic DNA was extracted from the legs and wings of the adult male moth using Qiagen G-tips. Briefly, we pulverized 14mg of wing and leg tissue to a fine powder with a RETSCH CryoMill. DNA was purified from this powder using the Qiagen Genomic-tip 20/G kit with a modified lysis buffer consisting of 20mM EDTA, 100mM NaCl, 1% Triton-X, 33mM Guanidine Thiocyanate and 10mM Tris-HCl. The 50C lysis step was extended overnight. The DNA quantity and quality were measured with Qubit 3.0 (Thermo Fisher Scientific, Inc., Carlsbad, CA, USA). High-quality DNA was used for library preparation and high throughput sequencing with Oxford Nanopore and Illumina platforms (Table 1).

**Table 1:** Sequencing Summary Statistics

| Library Type | Platform | Yield (Gb) | Library Size (bp) | Number of reads | Applications |
| --- | --- | --- | --- | --- | --- |
| Short read | Illumina Novaseq | 32.83 | 2 X 150 | 219M | Correction |
| Long read | ONT MinION | 19.47 | N50: 9,156 | 4.40M | Assembly |
| Hi-C | Illumina Novaseq | 32.76 | 2 X 150 | 218M | Scaffolding |

Oxford nanopore sequencing libraries were prepared using the Ligation Sequencing 1D Kit (Oxford Nanopore, Oxford, UK, p/n SQK-LSK109) according to manufacturer's instructions and sequenced for 48 hours on three MinION R9.4.1 flow cells. Nanopore reads were base-called with ONT Albacore Sequencing Pipeline Software (version 2.1.10). Reads from three flow cells were combined, and microbial read sequences identified by Centrifuge (v1.0.3-beta) were removed from Oxford Nanopore data [11]. This resulted in a total of 4,404,206 reads with a read length N50 of 9,156 bp and a total of 19.5 Gb (Table 1; Supplementary Table S1 and Supplementary Figure S1). For shotgun Illumina sequencing, a paired-end (PE) library was prepared with the Nextera DNA Flex Library Prep Kit from Illumina and sequenced on the Illumina NovaSeq6000 (Illumina, Inc., San Diego, CA, USA) yielding 219M paired-end 150 (PE150) reads. A total of 32.83 Gb of Illumina data were generated and used for genome survey, correction, and evaluation. All sequencing data have been deposited at the NCBI SRA database under BioProject PRJNA658700.

### Hi-C

To assemble contigs into chromosome sized scaffolds, we generated Hi-C libraries using 200mg of tissue from the head of the same animal. The tissue was ground with a mortar and pestle cooled by liquid nitrogen. We then followed the Phase Genomics Proximo HiC animal kit (Phase Genomics, Inc., Seattle, WA, USA) protocol. The quality of the resulting purified library was evaluated with Qubit 3.0 (Thermo Fisher Scientific, Inc.), and the Agilent 4200 TapeStation System (Agilent Technologies, Inc., Santa Clara, CA, USA). For a final QC step, we ran the library on a 2×100 cycle sequencing run on an Illumina MiSeq v2 (Illumina, Inc., San Diego, CA, USA). The qualified library was sequenced using the Illumina NovaSeq6000 (Illumina, Inc., San Diego, CA, USA) again generating 150bp paired-end reads. A total of 218M reads (32.76 Gb) were generated on the NovaSeq6000 and used for the subsequent Hi-C analysis (NCBI SRA BioProject PRJNA658700).

### Genome Assembly and Polishing

Nanopore sequencing reads were used to construct an initial assembly using Canu (v2.0) [12]. This initial assembly contained 5,381 contigs with a contig N50 of 424,554bp. Next, we polished the Canu assembly by first aligning all nanopore data to the draft assembly with Minimap2 (2.17-r943-dirty) and running a single iteration of the nanopolish (v0.11.1) consensus module [13,14]. Nanopolish uses a hidden Markov model (HMM) to examine the raw electrical data for possible improvements to the consensus sequence. The polished nanopore draft assembly was then error-corrected with 32.38 Gb of shotgun Illumina data using Bowtie2 (v2.4.1) for alignment and Racon (v1.3.3) for polishing [15,16]. We iteratively polished the Canu assembly with Racon, and after each iteration, the polished genome was aligned to the previous genome and SNPs were called with Mummer4 to evaluate the number of bases changed [17]. To assess the improvements made by polishing we ran BUSCO, a quantitative assessment of genome assembly and completeness based on evolutionarily informed expectations of gene content [18]. Errors in the assembly cause genes to go undetected by BUSCO, therefore being labeled as “missing” or “fragmented” BUSCOs. A single iteration of nanopolish improved BUSCO completeness scores by 4.0% (Figure 1B). A single iteration of Racon improved BUSCO completeness scores by 19.6% (Figure 1B) and further iterations had minimal improvements (Supplementary Figure S3).

**Figure 1:**
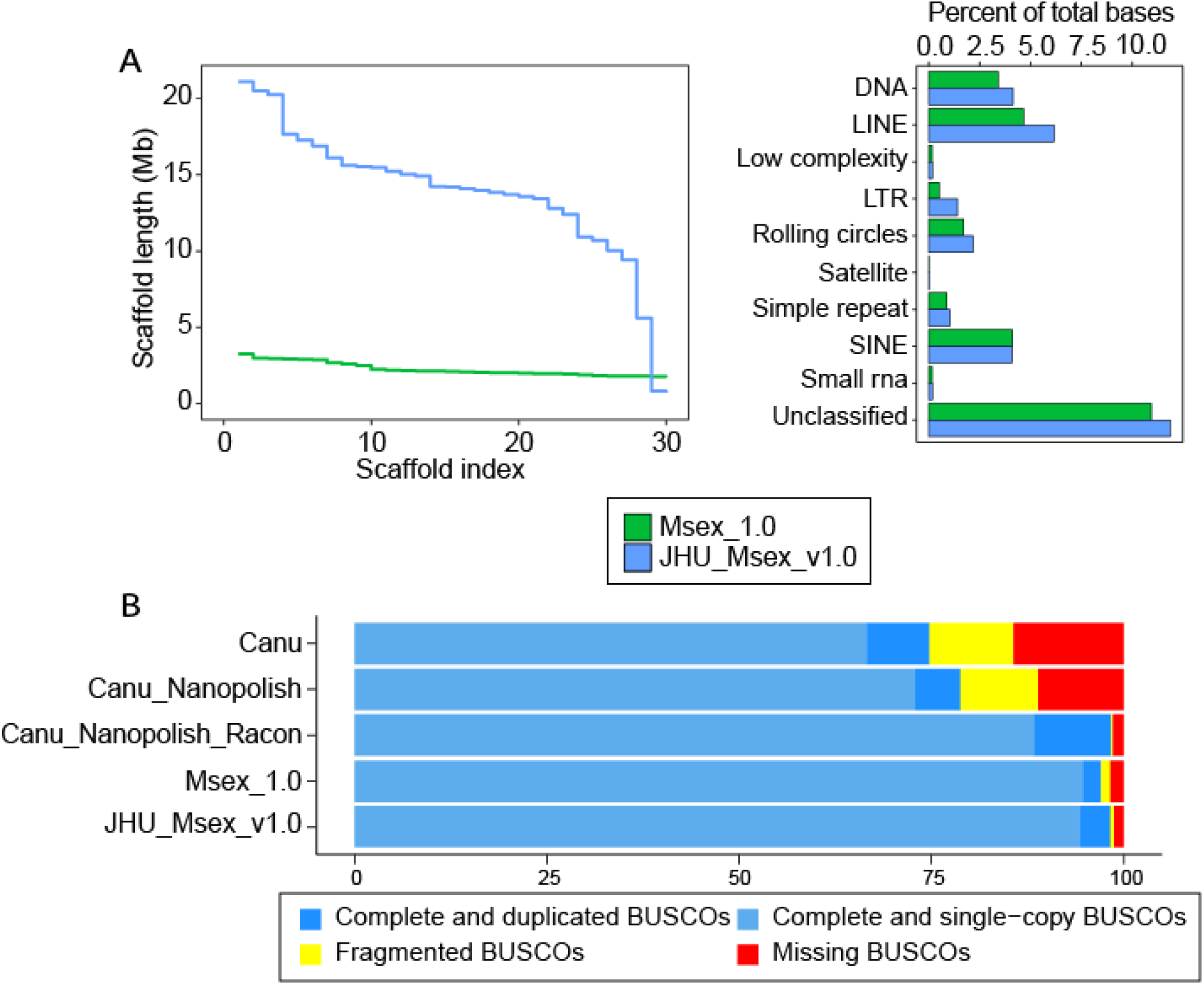
Final assembly metrics. **A)** (Left) NGx plot for final JHU_Msex_v1.0 assembly after scaffolding compared to the old Msex_1.0 assembly. Plot represents the largest 30 scaffolds from each assembly. (Right) Repeat annotation comparisons JHU_Msex_v1.0 compared to MSex_1.0. **B)** BUSCO results from insecta odbv10 database. Comparing the raw Canu assembly, Nanopolished Canu assembly, Racon and Nanopolished Canu assembly, the final scaffolded and polished assembly (JHU_Msex_v1.0) and the old Msex_1.0 assembly.

### Chromosome Assembly

To improve alignment of HiC libraries we trimmed 150 bp reads to 75 bp with TrimGalore (v0.6.0) (--hardtrim5 75) (http://www.bioinformatics.babraham.ac.uk/projects/trim_galore/). To connect the polished contigs into chromosome-scale scaffolds, we used the Hi-C data with the 3D-DNA (v180922) and Juicer (v1.6) pipelines (--mode diploid --editor-repeat-coverage 3) [19]. 3D-DNA generated 4,057 scaffolds of which 28 contain 86% (404.70 Mb) of the total input sequence length, consistent with the 28 chromosomes seen on previous karyotype analyses (Supplementary Figure S2) [20]. The final chromosome level assembly of *M. sexta* is 470 Mb (467 Mb without gaps) with a contig N50 of 402 Kb and a scaffold N50 of 14.2 Mb, a considerable improvement when compared to Msex_1.0 (Table 2, Figure 1A). We evaluated the final assembly for completeness with BUSCO insecta_odb10 (Figure 1B; Supplementary Table S2). We note that our final scaffolded *M. sexta* assembly is highly complete and contiguous, containing 98.1% complete insecta BUSCOs (Figure 1B).

**Table 2:** Final assembly statistics

|  | JHU_Msex_V1.0 | Msex_1.0 |
| --- | --- | --- |
| Total contig length | 468,966,500 | 399,658,702 |
| Number of contigs | 6,517 | 38,553 |
| Contig N50 | 402,416 | 40,289 |
| Largest contig | 2,620,325 | 401,033 |
| Total scaffold length | 470,051,811 | 419,427,777 |
| Number of scaffolds | 4,057 | 20,869 |
| Scaffold N50 | 14,248,853 | 664,006 |
| Largest scaffold | 21,114,789 | 3,253,989 |

### Repetitive elements

With the genome in hand, we next examined the repetitive elements in the *M. sexta* genome assembly. We used RepeatModeler2 to generate *de novo* repeat libraries [21]. Next, we performed homology based repeat masking using the *de novo* libraries in combination with the curated Metazoan library with RepeatMasker. We detected 159 Mb of repetitive sequence representing 33.91% of the genome, which is greater than the 28.84% detected from the Msex_1.0 assembly (Figure 1A, Supplementary Table S3). This proportion of repetitive sequence is considerably smaller than that of the Domestic silkworm genome (*Bombyx mori)* (~46.84% of 460 Mb genome) [8], but notably greater than reported for the monarch butterfly, *D. plexippus* (~13% of 273Mb) [22]. Among classified repeats, LINE elements were the most abundant superfamily found in the *M. sexta* genome. However, the overwhelming majority of masked regions dispersed throughout the genome corresponded to complex repetitive sequences yet to be characterized (11.88%), which is consistent with the Domestic silkworm (11.73%).

### Gene annotation

Gene annotation was primarily accomplished via a liftover of the Msex_1.0 annotations from the NCBI annotation (GCA_000262585.1) with Liftoff, a tool that maps annotations between closely related species or, in this case, two assemblies of the same species [23]. Liftoff successfully mapped 39,957 of the 41,110 original features to our new *M. sexta* assembly. We supplemented these annotations with gene models assembled from RNA-seq data and *ab initio* gene predictions made by AUGUSTUS [24]. We collected 67 publicly available *M. sexta* RNA-seq datasets collected from a variety of tissue and life stages (Supplementary Data 1) [25]. All RNA-seq sequences were first trimmed with TRIMMOMATIC (SLIDINGWINDOW:4:20 LEADING:10 TRAILING:10 MINLEN:50) prior to alignment to the *M. sexta* assembly with HISAT2 using default parameters [26,27]. An average of 96.2% of reads aligned from all libraries (Supplementary Data 1). This data was input into the BRAKER pipeline which relies on GenMark-ES/ET and AUGUSTUS to generate gene predictions [28–35]. Additionally, we assembled the full transcriptome from the RNA-seq data with Stringtie2 [36]. Both the de novo transcriptome assembly and *ab initio* gene predictions are prone to assembling erroneous transcripts. In order to only filter for high quality transcripts, we first aligned both the Stringtie2 and AUGUSTUS gene models to the Liftoff gene models with GFFCompare [37]. To identify unannotated genes, we examined transcripts supported by evidence from both the Stringtie2 assembly and the AUGUSTUS prediction, but not contained in the Liftoff annotation. We identified 794 such transcripts to add to the annotation. Overall, we identified 25,256 total genes compared to the 24,462 contained in the NCBI annotation of Msex_1.0.

### Identification of orthologous genes and phylogenetic tree construction

To identify candidate coding regions and generate predicted protein sequences, we ran TransDecoder (https://github.com/TransDecoder/TransDecoder/wiki) on our annotation. The generated protein sequences were used for determining phylogenetic relationships between *M. sexta* and other lepidopteran species by measuring pairwise sequence similarity. We used OrthoFinder (v2.3.12) to identify orthologous gene clusters in *M. sexta* and five other related lepidopteran species: *Bombyx mori* (domestic silkworm, GCF_000151625.1), *Plutella xylostella* (diamondback moth, GCF_000330985.1), *Papilio polytes* (common Mormon, GCF_000836215.1), *Papilio xuthus* (Asian swallowtail, GCF_000836235.1), and *Danaus plexippus plexippus* (monarch butterfly, GCF_009731565.1)[32,38–48]. OrthoFinder groups genes into orthogroups, sets of genes descended from a single gene in the species last common ancestor based on their sequence similarity.

OrthoFinder identified 15,428 orthogroups containing 120,091 total genes. Of these, 8,239 (53.4%) were shared between all six species and 1,783 were shared and single copy (Figure 2A; Supplementary Figure S4). Furthermore, the Tobacco hornworm had the most shared orthogroups with the Domestic silkworm (Figure 2A). The phylogenetic tree indicated the Tobacco hornworm is most closely related to the Domestic silkworm (Figure 2B). This closer relationship is expected due to the fact that both are members of the Lepidopteran superfamily Bombycoidea [49].

**Figure 2:**
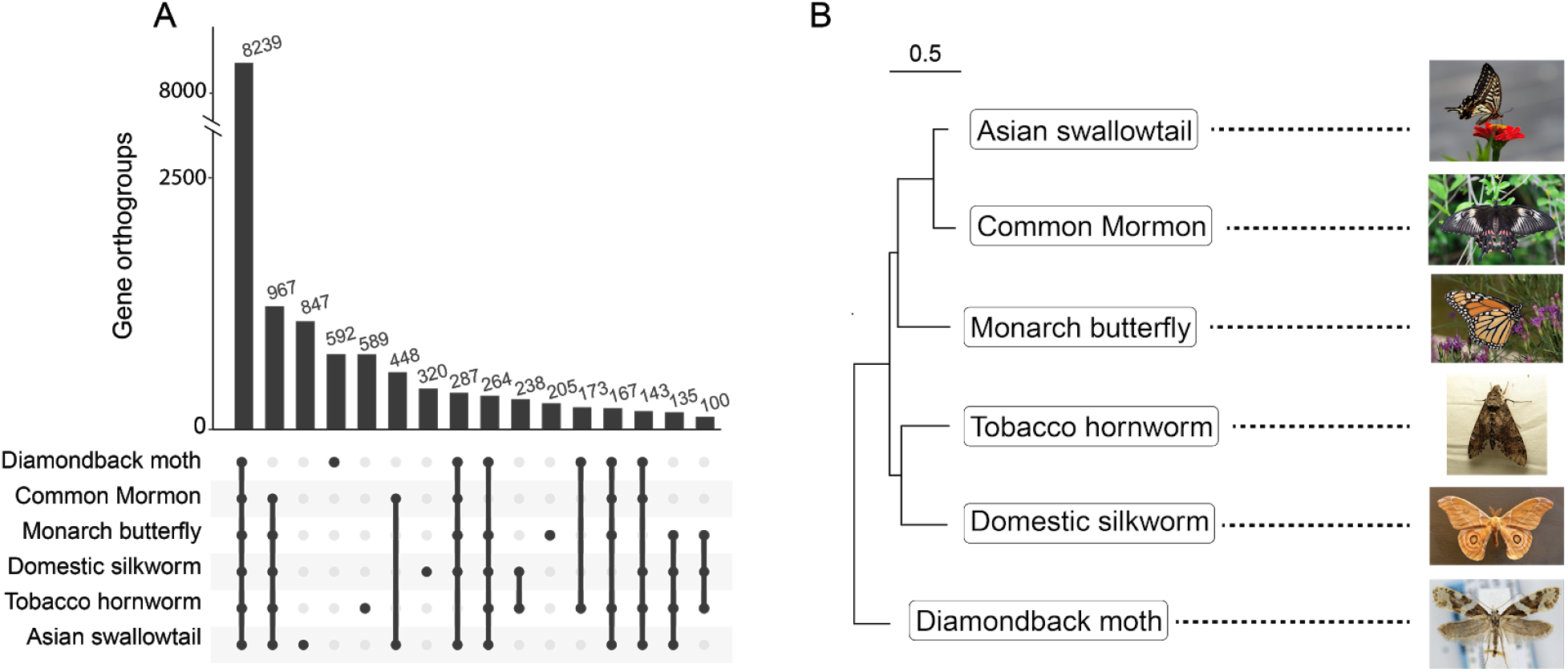
Phylogenetic relationships. **A)** Upset plot illustrating the number of shared orthogroups between the six species. Bars with less than 100 orthogroups were removed. **B)** Phylogenetic tree generated from orthogroup comparisons.

### Expression analysis

Using the same RNA-seq data for annotation, we wanted to examine how our assembly improved results of gene expression analysis. The average alignment rate from each dataset to JHU_Msex_v1.0 was 96.16% and to the Msex_1.0 assembly was 88.23% (Figure 3; Supplementary Data 1). We noted that paired-end data sets aligned significantly better to our assembly than to the old assembly with an increase of 11.92% of reads aligned as compared to single-end libraries where we only saw an improvement 4.06% of reads aligning. Gene abundance values were calculated using our annotation with the Stringtie2 gene abundance pipeline and the regularized log transformation (rlog) from DEseq2 [50]. The rlog transformation takes the read count data from Stringtie2 and accounts for differences in sequencing depth, RNA composition, heteroskedasticity, and large dynamic range [50]. To filter out genes with low expression in all tissues, we only retained genes if they had an rlog score of greater than three in at least one library. We then calculated the z-score of the rlog values and computed a euclidean distance matrix to perform hierarchical clustering which resulted in 18 gene clusters (Figure 3). To annotate the gene clusters we ran Interproscan5 to add GO annotations for all genes then used TopGO for GO term enrichment calculations within each gene cluster [51,52]. Significantly enriched GO terms for biological process (BP), molecular function (MF) and cellular compartment (CC) were calculated by Fisher's exact test and determined significant if the p-value was less than .05. We note that these RNA-seq datasets were collected from different animals without biological replicates and throughout multiple RNA-seq studies and while results show interesting patterns, we cannot make definitive biological conclusions.

**Figure 3:**
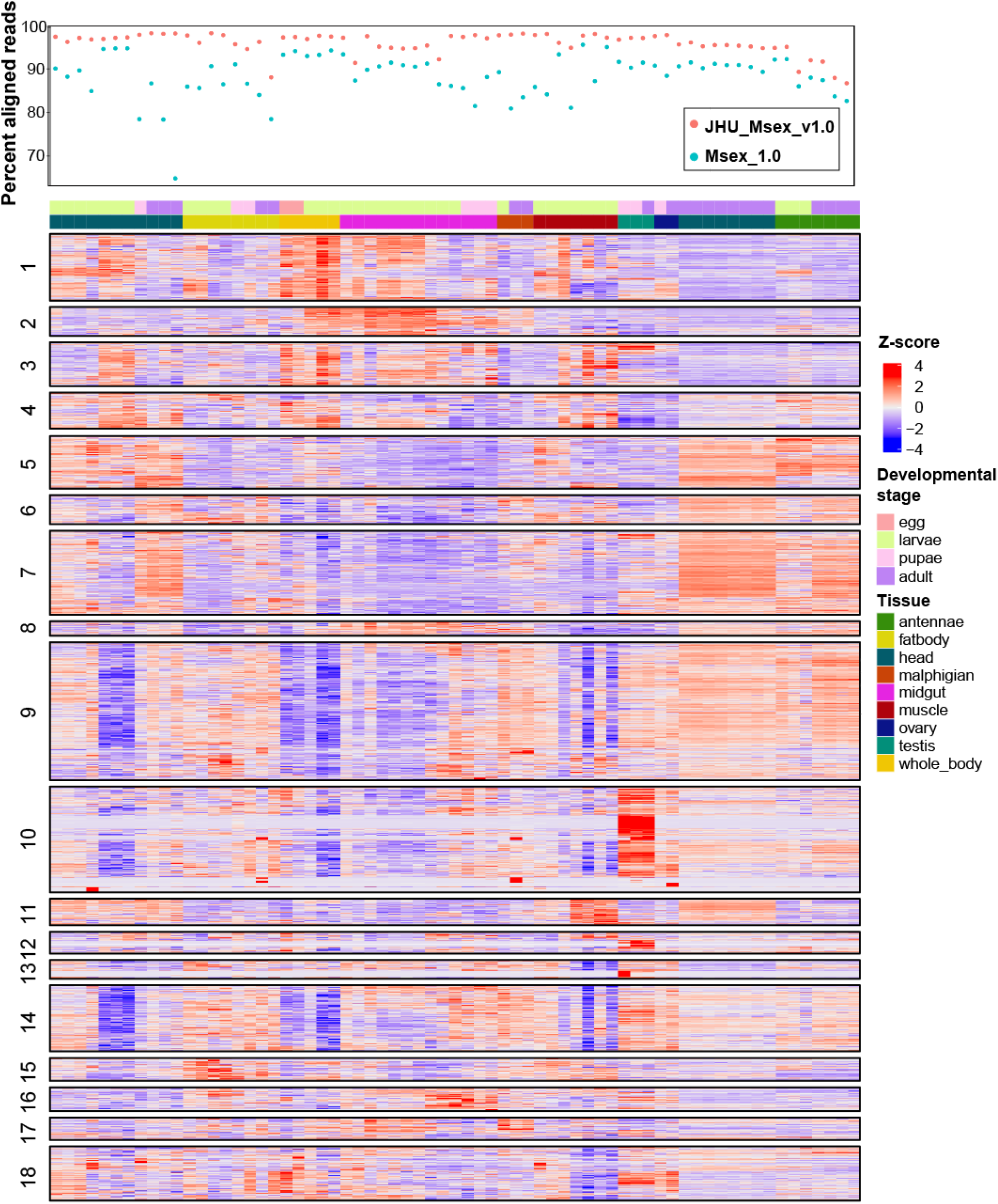
Gene expression clustering. Top panel is the percent aligned from each RNA-seq library to both our assembly and the old assembly. Expression matrix is the Z-score of the rlog expression transformation for all highly expressed genes. Genes were clustered by Z-score into 18 clusters by euclidean distance clustering. Libraries are in order shown in Supplementary data 1.

As expected, the enriched GO terms were well correlated with the expression pattern of gene clusters. For example, cluster 10 has the highest expression in the ovaries and testes and the most significantly enriched terms of cluster 10 include microtubule-based process, lncRNA processing and transcription (Supplementary Data 2). Sperm generation and meiosis requires continuous cell division and chromosome segregation, aided by the movement of microtubules. Additionally, the flagellar motor of sperm cells that aids in their motility is made up of microtubules. Cluster 7 includes genes highly expressed in the antennae and adult head. Consistent with these tissues being responsible for sensory perception, GO annotations are enriched for GTPase activity and odorant binding (Supplementary Data 2). Insects have a repertoire of sensory driven behaviors and odorant receptor (ORs) genes code for an entire family of G protein coupled receptors (GPCRs), a class of transmembrane proteins that have GTPase activity [53].

We also noted that both clusters 2 and 16 were highly expressed in the midgut tissue, albeit at different developmental stages. Cluster 2 contains genes involved in proteolysis, metabolic processes and catalytic activity and the expression is high in the larval stages of the midgut until larval stage five (L5), the pre-wandering stage. In the stages after L5, expression of cluster 16 genes increases in the midgut (Supplementary Data 2). Cluster 16 contains genes involved in the negative regulation of metabolic processes and apoptotic processes (Supplementary Data 2). During the process of metamorphosis the moth larvae will stop feeding in the pre-wandering stage, which coincides with the decrease in expression of the cluster 2 genes and increase in the negative regulation of metabolic processes. During the pupation process the insects experience massive tissue reorganization including death of major organs [54,55]. This process is likely mediated by apoptotic or autophagic genes, eg. Autophagy-related protein 8 (ATG8), as seen with *B. mori [56]*.

### Metamorphosis in the midgut

To showcase our new reference genome, we focused on gene expression changes occurring in the midgut during *M. sexta* development. We focused on serine proteases, which are proteins involved in catalytic cleavage of other proteins, significant in physiological processes like digestion, development, and defense. We ran Interproscan5 to identify putative serine proteases by Pfam classification [57]. We identified 240 proteins with potential serine protease domains (PF00089), compared to the 193 identified in Msex_1.0 [10]. Of the serine proteases we identified 76 that were highly expressed primarily in the midgut, compared to the 68 gut proteases previously identified in Msex_1.0 [10](Figure 4A). Upon the cessation of feeding in the L5 pre-wandering stage, expression of the digestive proteases declines rapidly as the animals prepare for pupation (Figure 4A,B). With our new highly contiguous genome assembly we were able to annotate large protease gene arrays on scaffolds 13 (60kb) and 18 (1Mb).

**Figure 4:**
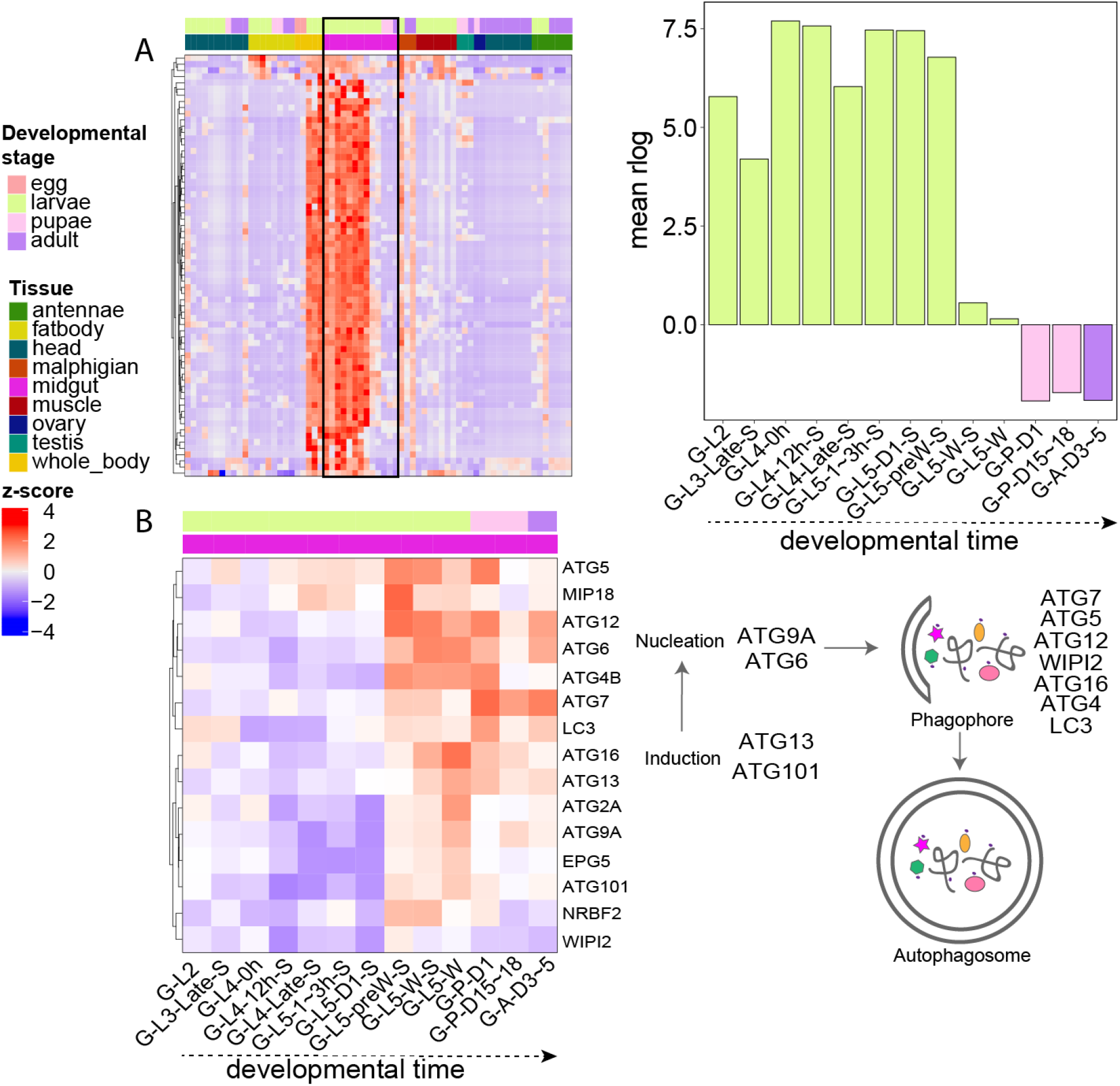
Gene expression in the midgut. **A)** (Left) Heatmap of Z-scores for expression of digestive proteases. (Right) Quantification of midgut digestive protease expression throughout development. Library naming nomenclature was derived from Cao & Jiang 2017 [25]. The first part of the library names indicates that the libraries are made from midgut (G). The second part indicates major stages of the insect, i.e. embryo (E), 1st to 5th instar larvae (L1 − L5), pupae (P), and adults (A). In the third part, “D” stands for day, “h” for hour, “preW” for pre-wandering, “W” for wandering. “S” in the last part of library names indicates single-end sequencing; no “S” in the end indicates paired-end sequencing. The libraries present as follows: midgut (G) (2nd L; 3rd L; 4th L, 0 h; 4th L, 12 h; 4th L, late; 5th L, 1–3 h; 5th L, 24 h;. 5th L, preW; 5th L, W; P, D1; P, D15–18; A, D3–5) **B)** (Left) Heatmap of expression Z-score of midgut autophagic genes throughout development. Gene names were assigned based on the NCBI GCA_000262585.1. Genes not present in this annotation were functionally annotated with Interproscan5 and assigned gene names. All genes in the heatmap were annotated as autophagy Gene Ontology Term (GO:0006914). (Right) Overview of the autophagic pathway [63].

*M. sexta* metamorphosis represents a critical developmental phase where the larval body is reorganized and larval organs can completely degenerate due autophagic and apoptotic cell death processes. The term “autophagic cell death” was introduced while studying metamorphic degeneration of muscle tissue in silkmoths [58]. Since then, the autophagic pathway has been studied extensively during the metamorphosis of many other lepidopterans. For example, in fruit flies, autophagy was demonstrated to be required for the removal of salivary glands and the midgut [59], however, apoptosis was shown to drive abdominal muscle remodelling [60]). To further investigate the mechanism of midgut tissue reorganization during metamorphosis we analyzed the gene expression patterns of genes involved in both apoptosis and autophagy in the *M. sexta* midgut throughout development. We generated lists of autophagy and apoptosis related genes using the GO annotations GO0006914 and GO0006915 from Interproscan5 as those encompassed multiple Pfam domains and gave a more complete list of genes related to these pathways. We did not note a distinct expression pattern of apoptotic genes throughout midgut development (Supplementary Figure S5). However, we did identify an increase in expression of genes within the autophagic pathway beginning at the L5 pre-wandering stage (G-L5-preW-S), which coincides with the decline in digestive protease expression at the L5 wandering stage (G-L5-W-S/G-L5-W) (Figure 4B). These results indicate the potential involvement of the autophagic pathway in *M. sexta* midgut during metamorphosis, at the same time in development that autophagy has been shown to remodel the midgut of other insects such fruit flies [61], silkmoths [56], and sand flies [62].

### Reuse potential

Our results have leveraged short and long read sequencing technology to assemble a highly contiguous reference genome of *M. sexta*. We lifted over the original genome annotations to this improved assembly, maintaining functional annotations and gene IDs for researchers currently studying *M. sexta* genes. We believe the improved assembly will aid current and future studies using *M. sexta* as a model system for research on fundamental processes in insect physiology and biochemistry.

## Data availability

The JHU_Msex_v1.0 genome and corresponding annotation is available on Zenodo (10.5281/zenodo.4005068). Whole genome sequencing data are available in the NCBI SRA database under BioProject PRJNA658700. The scripts we used in this article, including the genome assembly, genome polishing, repeat annotation, and genome assessments, are available on Github (https://github.com/timplab/moth).

## Acknowledgements

This work was partially supported by NIH Ruth L. Kirschstein Institutional National Research Service Award (T32) GM007445.

## References

1. Wieczorek H, Grüber G, Harvey WR, Huss M, Merzendorfer H. The plasma membrane H+-V-ATPase from tobacco hornworm midgut. J Bioenerg Biomembr. 1999;31:67–74.

2. Kay I, Patel M, Coast GM, Totty NF, Mallet AI, Goldsworthy GJ. Isolation, characterization and biological activity of a CRF-related diuretic peptide from Periplaneta americana L. Regul Pept. 1992;42:111–22.

3. Truman JW, Reiss SE. Neuromuscular metamorphosis in the moth Manduca sexta: hormonal regulation of synapses loss and remodeling. J Neurosci. 1995;15:4815–26.

4. Mechaber WL, Capaldo CT, Hildebrand JG. Behavioral responses of adult female tobacco hornworms, Manduca sexta, to hostplant volatiles change with age and mating status [Internet]. Journal of Insect Science. 2002. Available from: http://dx.doi.org/10.1093/jis/2.1.5

5. Kanost MR, Jiang H, Yu X-Q. Innate immune responses of a lepidopteran insect, Manduca sexta. Immunol Rev. 2004;198:97–105.

6. Lyons N, Softley I, Balfour A, Williamson C, O’Brien HE, Shetty AC, et al. Tobacco Hornworm (Manduca sexta) caterpillars as a novel host model for the study of fungal virulence and drug efficacy [Internet]. 2020 [cited 2020 Aug 14]. p. 693226. Available from: https://www.biorxiv.org/content/10.1101/693226v2.abstract

7. de Boer G, Hanson FE. Feeding responses to solanaceous allelochemicals by larvae of the tobacco hornworm, Manduca sexta. Entomol Exp Appl. Wiley Online Library; 1987;45:123–31.

8. Kawamoto M, Jouraku A, Toyoda A, Yokoi K, Minakuchi Y, Katsuma S,et al. High-quality genome assembly of the silkworm, Bombyx mori. Insect Biochem Mol Biol. 2019;107:53–62.

9. Gu L, Reilly PF, Lewis JJ, Reed RD, Andolfatto P, Walters JR. Dichotomy of Dosage Compensation along the Neo Z Chromosome of the Monarch Butterfly. Curr Biol. 2019;29:4071–7.e3.

10. Kanost MR, Arrese EL, Cao X, Chen Y-R, Chellapilla S, Goldsmith MR, et al. Multifaceted biological insights from a draft genome sequence of the tobacco hornworm moth, Manduca sexta. Insect Biochem Mol Biol. 2016;76:118–47.

11. Kim D, Song L, Breitwieser FP, Salzberg SL. Centrifuge: rapid and sensitive classification of metagenomic sequences. Genome Res. 2016;26:1721–9.

12. Koren S, Walenz BP, Berlin K, Miller JR, Bergman NH, Phillippy AM. Canu: scalable and accurate long-read assembly via adaptive k-mer weighting and repeat separation. Genome Res. 2017;27:722–36.

13. Li H. Minimap2: pairwise alignment for nucleotide sequences. Bioinformatics. 2018;34:3094–100.

14. Loman NJ, Quick J, Simpson JT. A complete bacterial genome assembled de novo using only nanopore sequencing data. Nat Methods. 2015;12:733–5.

15. Vaser R, Sović I, Nagarajan N, Šikić M. Fast and accurate de novo genome assembly from long uncorrected reads. Genome Res. 2017;27:737–46.

16. Langmead B, Salzberg SL. Fast gapped-read alignment with Bowtie 2. Nat Methods. 2012;9:357–9.

17. Marçais G, Delcher AL, Phillippy AM, Coston R, Salzberg SL, Zimin A. MUMmer4: A fast and versatile genome alignment system. PLoS Comput Biol. 2018;14:e1005944.

18. Simão FA, Waterhouse RM, Ioannidis P, Kriventseva EV, Zdobnov EM. BUSCO: assessing genome assembly and annotation completeness with single-copy orthologs. Bioinformatics. 2015;31:3210–2.

19. Dudchenko O, Batra SS, Omer AD, Nyquist SK, Hoeger M, Durand NC, et al. De novo assembly of the Aedes aegypti genome using Hi-C yields chromosome-length scaffolds. Science. 2017;356:92–5.

20. Yasukochi Y, Tanaka-Okuyama M, Shibata F, Yoshido A, Marec F, Wu C, et al. Extensive conserved synteny of genes between the karyotypes of Manduca sexta and Bombyx mori revealed by BAC-FISH mapping. PLoS One. 2009;4:e7465.

21. Flynn JM, Hubley R, Goubert C, Rosen J, Clark AG, Feschotte C, et al. RepeatModeler2: automated genomic discovery of transposable element families [Internet]. Available from: http://dx.doi.org/10.1101/856591

22. Zhan S, Merlin C, Boore JL, Reppert SM. The monarch butterfly genome yields insights into long-distance migration. Cell. 2011;147:1171–85.

23. Shumate A, Salzberg SL. Liftoff: an accurate gene annotation mapping tool [Internet]. bioRxiv. 2020 [cited 2020 Jul 28]. p. 2020.06.24.169680. Available from: https://www.biorxiv.org/content/10.1101/2020.06.24.169680v1.abstract

24. Stanke M, Keller O, Gunduz I, Hayes A, Waack S, Morgenstern B. AUGUSTUS: ab initio prediction of alternative transcripts. Nucleic Acids Res. 2006;34:W435–9.

25. Cao X, Jiang H. An analysis of 67 RNA-seq datasets from various tissues at different stages of a model insect, Manduca sexta. BMC Genomics. 2017;18:796.

26. Bolger AM, Lohse M, Usadel B. Trimmomatic: a flexible trimmer for Illumina sequence data. Bioinformatics. 2014;30:2114–20.

27. Kim D, Paggi JM, Park C, Bennett C, Salzberg SL. Graph-based genome alignment and genotyping with HISAT2 and HISAT-genotype. Nat Biotechnol. 2019;37:907–15.

28. Hoff KJ, Lange S, Lomsadze A, Borodovsky M, Stanke M. BRAKER1: Unsupervised RNA-Seq-Based Genome Annotation with GeneMark-ET and AUGUSTUS: Table 1 [Internet]. Bioinformatics. 2016. p. 767–9. Available from: http://dx.doi.org/10.1093/bioinformatics/btv661

29. Hoff KJ, Lomsadze A, Borodovsky M, Stanke M. Whole-Genome Annotation with BRAKER. Methods Mol Biol. 2019;1962:65–95.

30. Li H, Handsaker B, Wysoker A, Fennell T, Ruan J, Homer N, et al. The Sequence Alignment/Map format and SAMtools. Bioinformatics. 2009;25:2078–9.

31. Barnett DW, Garrison EK, Quinlan AR. BamTools: a C++ API and toolkit for analyzing and managing BAM files. academic.oup.com; 2011; Available from: https://academic.oup.com/bioinformatics/article-abstract/27/12/1691/255399

32. Buchfink B, Xie C, Huson DH. Fast and sensitive protein alignment using DIAMOND. Nat Methods. 2015;12:59–60.

33. Stanke M, Diekhans M, Baertsch R, Haussler D. Using native and syntenically mapped cDNA alignments to improve de novo gene finding. Bioinformatics. 2008;24:637–44.

34. Stanke M, Schöffmann O, Morgenstern B, Waack S. Gene prediction in eukaryotes with a generalized hidden Markov model that uses hints from external sources. BMC Bioinformatics. 2006;7:62.

35. Gremme G, Steinbiss S, Kurtz S. GenomeTools: a comprehensive software library for efficient processing of structured genome annotations. IEEE/ACM Trans Comput Biol Bioinform. 2013;10:645–56.

36. Kovaka S, Zimin AV, Pertea GM, Razaghi R, Salzberg SL, Pertea M. Transcriptome assembly from long-read RNA-seq alignments with StringTie2. Genome Biol. 2019;20:278.

37. Pertea G, Pertea M. GFF Utilities: GffRead and GffCompare [Internet]. F1000Research. 2020. p. 304. Available from: http://dx.doi.org/10.12688/f1000research.23297.1

38. Bertone P, Stolc V, Royce TE, Rozowsky JS, Urban AE, Zhu X, et al. SF Altschul, W. Gish, W. Miller, EW Myers, and DJ Lipman. Basic local alignment search tool. J Mol Biol, 215: 403–410, 1990. M. Ariel, J. McCarrey, and H. Cedar. Methylation patterns of testis-specific genes. Proc Natl Acad Sci, 88: 2317–2321, 1991. JF Bateman, S. Freddi, G. Nattrass, and R. Savarirayan. Tissue-specific rna surveil. Proc Natl Acad Sci. 1991;88:2317–21.

39. Price MN, Dehal PS, Arkin AP. FastTree 2--approximately maximum-likelihood trees for large alignments. PLoS One. Public Library of Science; 2010;5:e9490.

40. Katoh K, Standley DM. MAFFT multiple sequence alignment software version 7: improvements in performance and usability. Mol Biol Evol. 2013;30:772–80.

41. Kelly S, Maini PK. DendroBLAST: approximate phylogenetic trees in the absence of multiple sequence alignments. PLoS One. 2013;8:e58537.

42. Lefort V, Desper R, Gascuel O. FastME 2.0: A Comprehensive, Accurate, and Fast Distance-Based Phylogeny Inference Program: Table 1 [Internet]. Molecular Biology and Evolution. 2015. p. 2798–800. Available from: http://dx.doi.org/10.1093/molbev/msv150

43. Huerta-Cepas J, Serra F, Bork P. ETE 3: Reconstruction, Analysis, and Visualization of Phylogenomic Data. Mol Biol Evol. 2016;33:1635–8.

44. Van Dongen S. Graph clustering by flow simulation PhD thesis. 2000. University of Utrecht, The Netherlands.

45. Emms DM, Kelly S. OrthoFinder: phylogenetic orthology inference for comparative genomics. Genome Biol. 2019;20:238.

46. Emms DM, Kelly S. OrthoFinder: solving fundamental biases in whole genome comparisons dramatically improves orthogroup inference accuracy. Genome Biol. 2015;16:157.

47. Emms DM, Kelly S. STRIDE: Species Tree Root Inference from Gene Duplication Events. Mol Biol Evol. 2017;34:3267–78.

48. Emms DM, Kelly S. STAG: Species Tree Inference from All Genes [Internet]. 2018 [cited 2020 Aug 11]. p. 267914. Available from: https://www.biorxiv.org/content/10.1101/267914v1.abstract

49. Kitching IJ, Rougerie R, Zwick A, Hamilton CA, St Laurent RA, Naumann S, et al. A global checklist of the Bombycoidea (Insecta: Lepidoptera). Biodivers Data J. 2018;e22236.

50. Love MI, Huber W, Anders S. Moderated estimation of fold change and dispersion for RNA-seq data with DESeq2. Genome Biol. 2014;15:550.

51. Jones P, Binns D, Chang H-Y, Fraser M, Li W, McAnulla C, et al. InterProScan 5: genome-scale protein function classification. Bioinformatics. 2014;30:1236–40.

52. Alexa A, Rahnenführer J, Lengauer T. Improved scoring of functional groups from gene expression data by decorrelating GO graph structure. Bioinformatics. 2006;22:1600–7.

53. Benton R. On the ORigin of smell: odorant receptors in insects. Cell Mol Life Sci. 2006;63:1579–85.

54. Dai JD, Gilbert LI. Programmed cell death of the prothoracic glands of Manduca sexta during pupal-adult metamorphosis. Insect Biochem Mol Biol. 1997;27:69–78.

55. Zakeri Z, Quaglino D, Latham T, Woo K, Lockshin RA. Programmed cell death in the tobacco hornworm, Manduca sexta: alteration in protein synthesis. Microsc Res Tech. 1996;34:192–201.

56. Romanelli D, Casartelli M, Cappellozza S, de Eguileor M, Tettamanti G. Roles and regulation of autophagy and apoptosis in the remodelling of the lepidopteran midgut epithelium during metamorphosis. Sci Rep. 2016;6:32939.

57. Finn RD, Bateman A, Clements J, Coggill P, Eberhardt RY, Eddy SR, et al. Pfam: the protein families database. Nucleic Acids Res. 2014;42:D222–30.

58. Lockshin RA, Williams CM. Programmed cell death—V. Cytolytic enzymes in relation to the breakdown of the intersegmental muscles of silkmoths [Internet]. Journal of Insect Physiology. 1965. p. 831–44. Available from: http://dx.doi.org/10.1016/0022-1910(65)90186-1

59. Berry DL, Baehrecke EH. Growth arrest and autophagy are required for salivary gland cell degradation in Drosophila. Cell. 2007;131:1137–48.

60. Zirin J, Cheng D, Dhanyasi N, Cho J, Dura J-M, Vijayraghavan K, et al. Ecdysone signaling at metamorphosis triggers apoptosis of Drosophila abdominal muscles. Dev Biol. 2013;383:275–84.

61. Denton D, Shravage B, Simin R, Mills K, Berry DL, Baehrecke EH, et al. Autophagy, not apoptosis, is essential for midgut cell death in Drosophila. Curr Biol. 2009;19:1741–6.

62. Malta J, Heerman M, Weng JL, Fernandes KM, Martins GF, Ramalho-Ortigão M. Midgut morphological changes and autophagy during metamorphosis in sand flies. Cell Tissue Res. 2017;368:513–29.

63. Levine B, Kroemer G. Autophagy in the pathogenesis of disease. Cell. 2008;132:27–42.

