## Supplementary Material for "*De novo* genome assembly of the Tobacco Hornworm moth (*Manduca sexta*)"

**TABLE OF CONTENTS**

**Supplementary Tables:**

**Supplementary Table S1:** Summary statistics of nanopore reads used for Canu assembly

**Supplementary Table S2:** BUSCO results for all versions of assembly

**Supplementary Table S3:** Repeat annotation of JHU_Msex_v1.0 and Msex_1.0

**Supplementary Figures:**

**Supplementary Figure S1:** Weighted histogram of read length distribution for nanopore reads

**Supplementary Figure S2:** Assembly contiguity assessments

**Supplementary Figure S3:** BUSCO scores throughout iterative polishing

**Supplementary Figure S4:** Phylogenetic relationships between *M. sexta* and other lepidoptera

**Supplementary Figure S5:** Expression of apoptotic genes throughout midgut development

**Supplementary Data:**

**Supplementary Data 1:** Summary of RNA-seq data including library descriptions, accessions and alignment statistics for alignment to JHU_Msex_v1.0 and Msex_1.0

**Supplementary Data 2:** Statistically significant Gene Ontology (GO) terms for each expression cluster.

**Supplementary Table S1:** Nanopore reads used for Canu assembly

| Statistic | Number of bases |
|---|---|
| Mean read length | 4,421 |
| Mean read quality | 8.8 |
| Median read length | 2,299 |
| Median read quality | 9.6 |
| Number of reads | 4,404,206 |
| Read length N50 | 9,156 |
| Total bases | 19,468,814,476 |

**Supplementary Table S2:** BUSCO assessment results

| | Complete | Complete and single copy | Complete and duplicated | Fragmented | Missing |
|---|---|---|---|---|---|
| Canu | 74.8 | 66.6 | 8.2 | 10.9 | 14.3 |
| Canu + Nanopolish | 78.8 | 72.9 | 5.9 | 10.1 | 11.1 |
| Canu + Nanopolish + Racon | 98.4 | 88.4 | 10.0 | 0.2 | 1.4 |
| JHU_Msex_v2.0 | 98.1 | 93.1 | 5.0 | 0.7 | 1.2 |
| Msex_1.0 | 97.1 | 94.8 | 2.3 | 1.2 | 1.7 |

**Supplementary Table S3:** Repeat annotation

| Repeat | Number of elements | Length | Percent of sequence |
|---|---|---|---|
| SINE | 138,977 | 19,210,993 | 4.09 |
| LINE | 166,753 | 28,925,639 | 6.15 |
| LTR | 132,31 | 6,611,509 | 1.41 |
| DNA | 112,268 | 19,490,103 | 4.15 |
| Unclassified | 350,797 | 55,860,647 | 11.88 |
| Total interspersed | | 130,098,891 | 27.68 |
| Small RNA | 7,901 | 853,329 | 0.18 |
| Satellites | 2,514 | 95,604 | 0.02 |
| Simple repeats | 109,241 | 4,898,120 | 1.04 |
| Low complexity | 17,972 | 831,316 | 0.18 |
| Bases masked | | 159,380,633 | 33.91 |

**Supplementary Figures:**

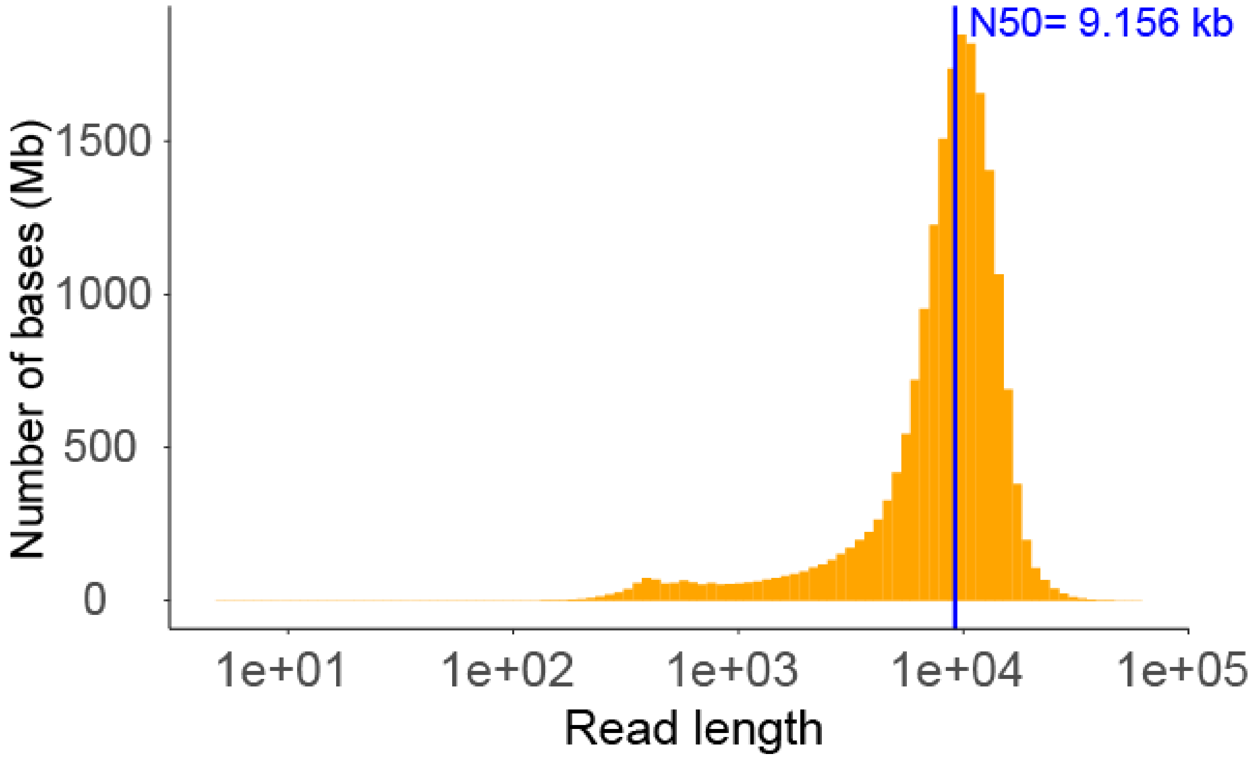


**Figure S1.** Weighted histogram of read length distribution for nanopore sequencing reads that were used in the Canu assembly.

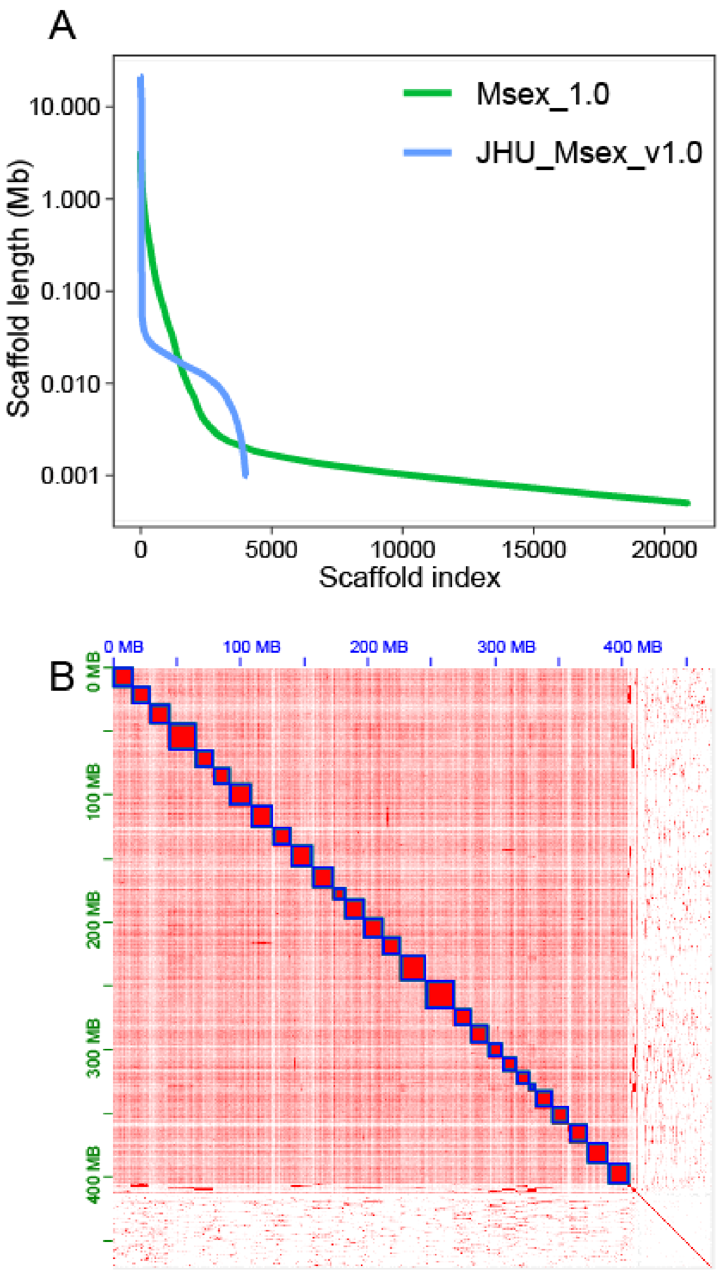


**Figure S2. A)** NGx plot of the entire assembly comparing JHU_Msex_v1.0 to Msex_1.0. **B)** HiC contact map generated by 3D-DNA and visualized in juicebox showing 28 chromosome size scaffolds (blue).

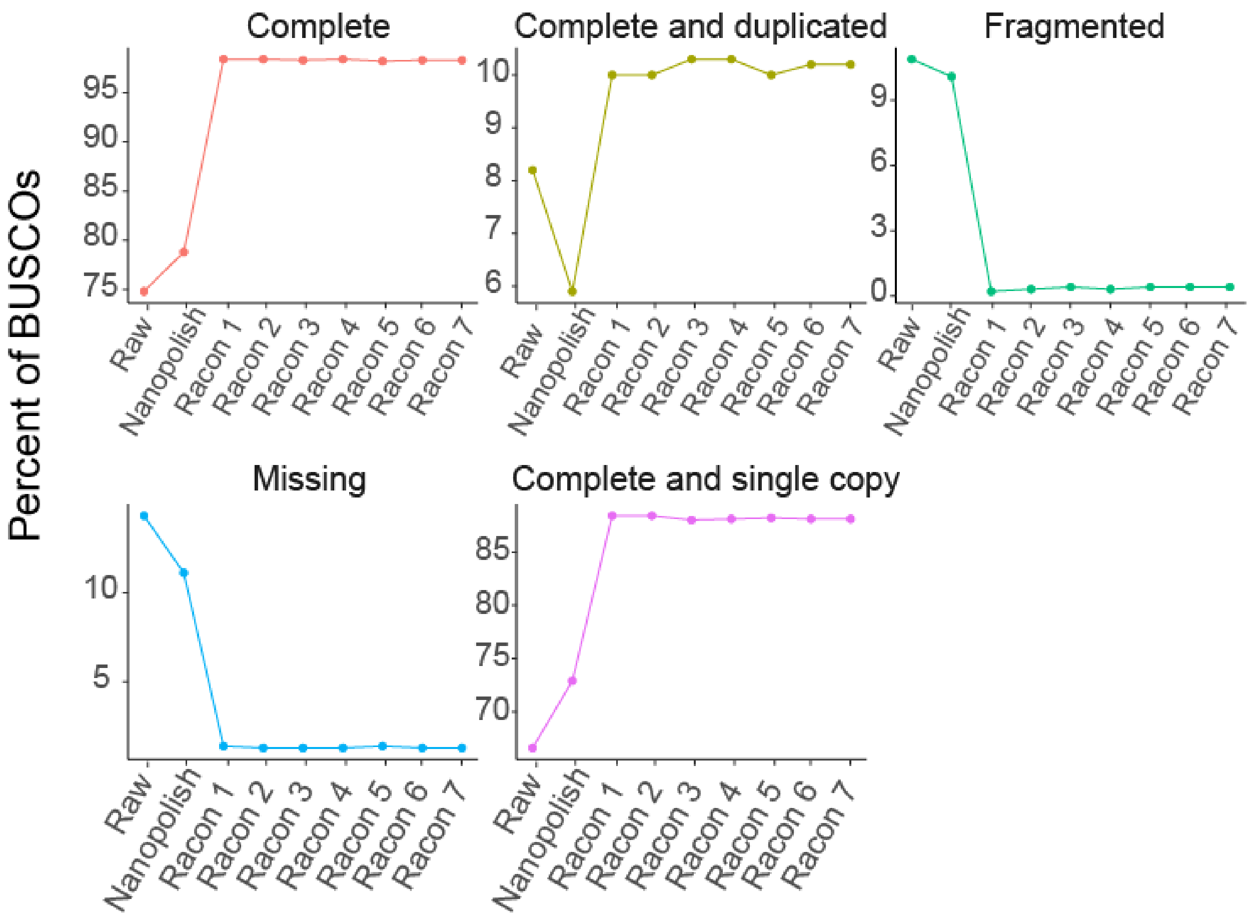


**Figure S3.** Determining number of polishing iterations using metrics from BUSCO insecta_odb10.

A

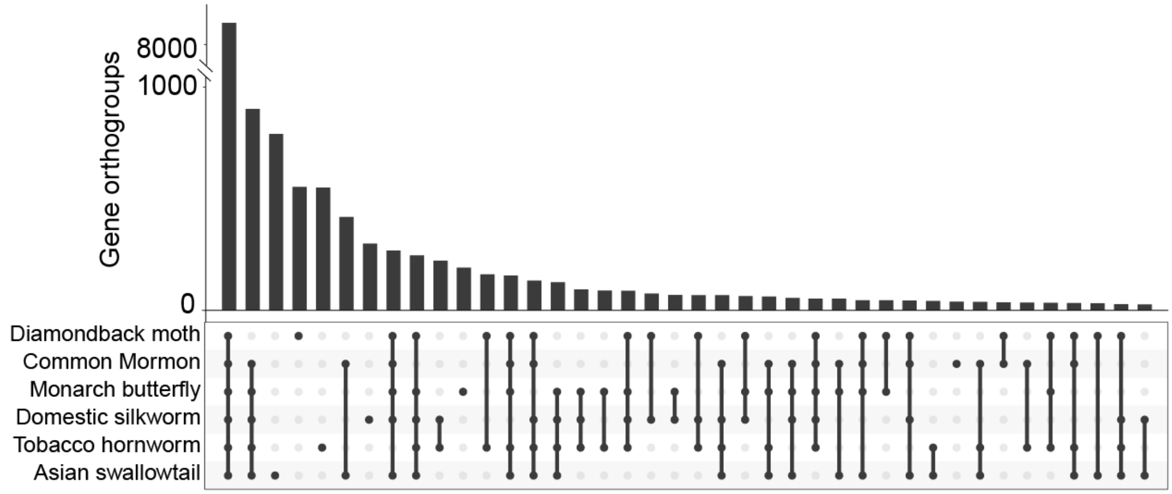


B

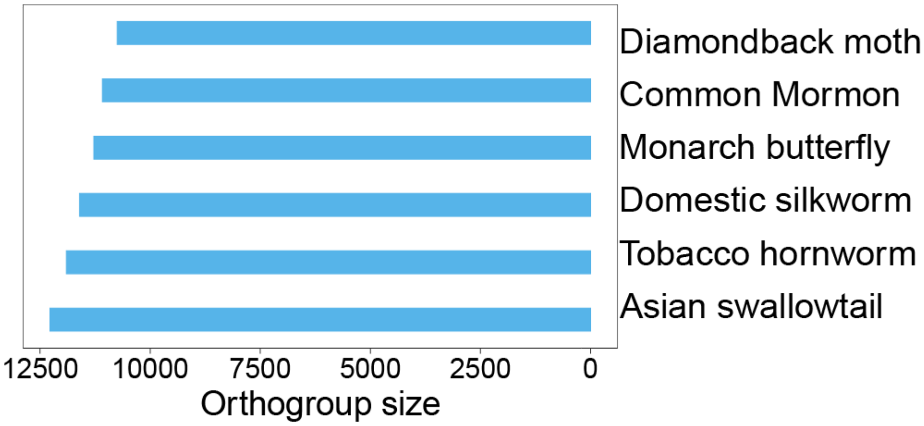


**Figure S4. Phylogenetic relationships A)** Upset plot illustrating the number of shared orthogroups between the six species. **B)** Number of gene orthogroups for each species.

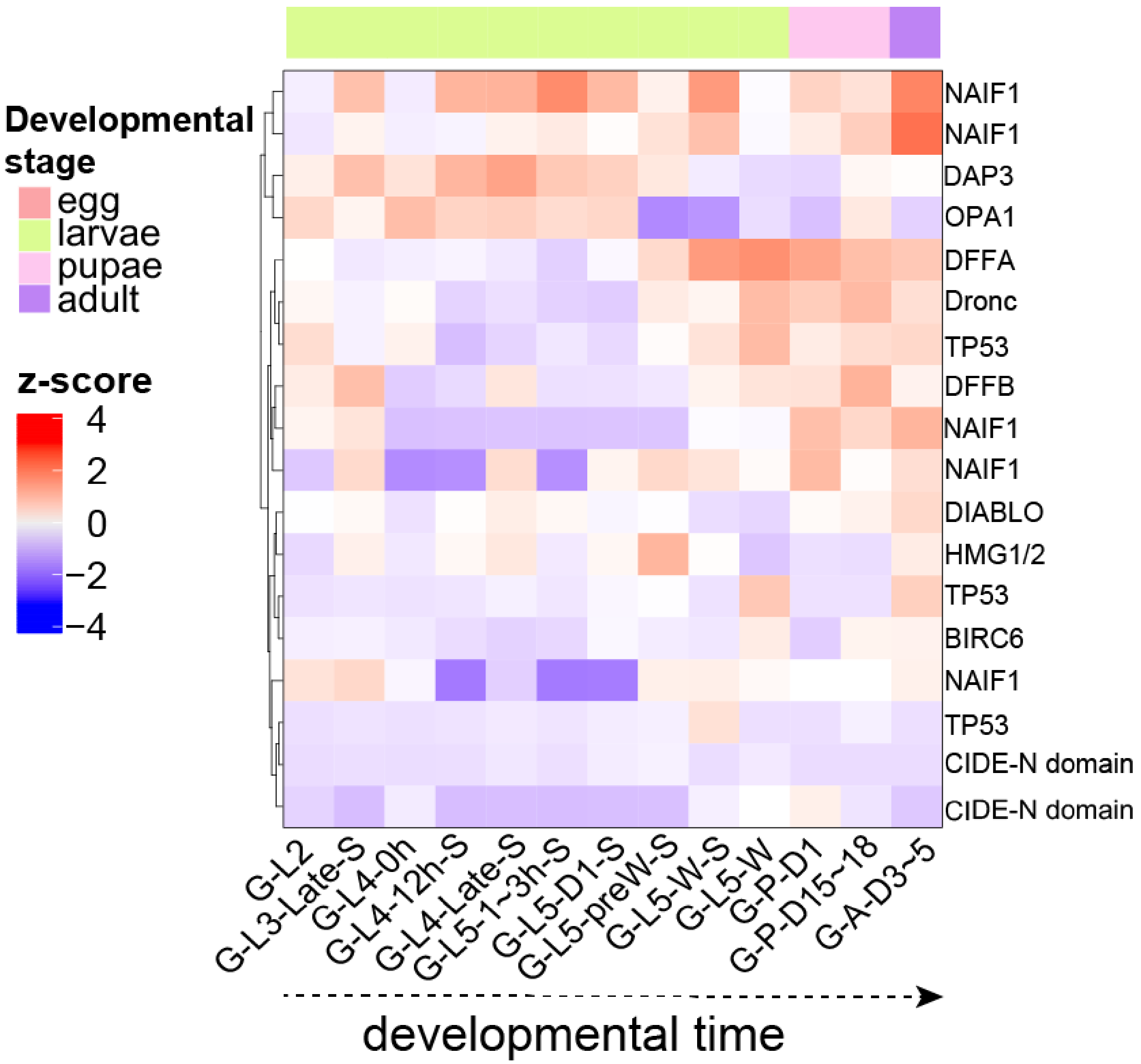


**Figure S5.** Expression of apoptotic genes in the midgut throughout developmental time. Genes were functionally annotated based on the NCBI GCA_000262585.1 gene functions. Genes not present in this annotation were functionally annotated with Interproscan5. All genes in the heatmap were annotated as apoptotic process Gene Ontology Term (GO:0006915).
